# Tactile sensory channels over-ruled by frequency decoding system that utilizes spike pattern regardless of receptor type

**DOI:** 10.1101/570929

**Authors:** Ingvars Birznieks, Sarah McIntyre, Hanna M Nilsson, Saad S Nagi, Vaughan G Macefield, David A Mahns, Richard M Vickery

**Affiliations:** School of Medical Sciences, Faculty of Medicine, UNSW Sydney, NSW 2052, Sydney, Australia; Neuroscience Research Australia, NSW 2031, Sydney, Australia; School of Medicine, Western Sydney University, NSW 2751, Sydney, Australia; The Baker Heart and Diabetes Institute. Victoria 3004, Melbourne, Australia; Biomedical Engineering and Neuroscience, MARCS institute, Western Sydney University, NSW 2214, Sydney, Australia; Linköping University, SE 581 83, Linköping, Sweden

**Keywords:** Tactile afferents, Pacinian channel, neural code, perception, vibrotactile stimuli, low threshold mechanoreceptors, hand function, spike timing, pitch

## Abstract

The established view is that vibrotactile stimuli evoke two qualitatively distinctive cutaneous sensations, flutter (frequencies < 60 Hz) and vibratory hum (frequencies > 60 Hz), subserved by two distinct receptor types (Meissner’s and Pacinian corpuscle, respectively) which may engage different neural processing pathways or channels and fulfill quite different biological roles. In psychological and physiological literature those two systems have been labelled as Pacinian and non-Pacinian channels. However, we present evidence that low-frequency spike trains in Pacinian afferents can readily induce a vibratory percept with the same low frequency attributes as sinusoidal stimuli of the same frequency thus demonstrating a universal frequency decoding system. We achieved this using brief low-amplitude pulsatile mechanical stimuli to selectively activate Pacinian afferents. This indicates that spiking pattern, regardless of receptor type, determines vibrotactile frequency perception. This mechanism may underlie the constancy of vibrotactile frequency perception across different skin regions innervated by distinct afferent types.

## Introduction

The sense of touch comprises a range of different perceptual qualities subserved by several distinct mechanoreceptor types and associated afferent nerve fibres in the skin (Johnson, 2001; Vallbo & Johansson, 1984). Observations that different receptor types are tuned to different stimulus features and have distinct response profiles have led researchers to conclude that different receptor types are the inputs to separate neural “channels” dedicated to processing of those features (Bolanowski, Gescheider, & Verrillo, 1994; Gescheider, 1976; Gescheider, Bolanowski, & Verrillo, 2004; Hyvarinen, Sakata, Talbot, & Mountcastle, 1968; Sretavan & Dykes, 1983). In the glabrous skin there are two types of fast adapting (FA) afferents which at the threshold level display characteristic U-shaped tuning curves to sinusoidal vibrotactile stimuli: FAI (or RA) afferents innervating Meissner’s corpuscles are preferentially activated at frequencies up to 60 Hz, while FAII (or Pacinian - PC) afferents innervating Pacinian corpuscles have much lower response thresholds, and are most sensitive to higher frequencies (> 100 Hz) (Talbot, Darian-Smith, Kornhuber, & Mountcastle, 1968). At the border between those two frequency domains, at about 60 Hz, there is a qualitative change in sensation from flutter to vibratory hum (Gescheider, 1976; LaMotte & Mountcastle, 1975; Talbot et al., 1968), which is used as further justification for psychophysical segregation into Pacinian and non-Pacinian channels. One assumption of this scheme is that the Pacinian channel does not possess neural circuits for processing low-frequency spiking patterns characteristic of low frequency sinusoidal stimuli and therefore cannot produce a perceptual experience outside the high frequency domain. This is based on laboratory testing using sinusoidal stimuli in which acceleration and periodicity are linked and thus Pacinian corpuscles wouldn’t respond at low frequencies. An intriguing question is the extent to which frequency processing circuitry is specialized for afferent type (FAI vs FAII) in their optimal sinusoidal frequency response range? There is a big gap in our knowledge as sinusoidal stimuli inherently don’t allow activation of FAII afferents at low frequencies and thus functionally are not representative for wide variety of natural stimuli involving discrete mechanical transients associated with motor control or surface structures with low spatial frequency.

We addressed this question by using brief pulsatile mechanical stimuli that enabled us to create arbitrary time-controlled spike trains of any frequency and pattern in the responding FAII afferents and thus investigate the perceptual properties of those spiking patterns (for details see (Birznieks & Vickery, 2017)). By setting the amplitude of the mechanical pulses below the FAI activation threshold, we first established that low frequency discharge in FAII afferents (the Pacinian channel) can indeed cause conscious perception of a tactile stimulus at frequencies as low as 6 Hz. We then investigated the perceptual properties of low frequency FAII afferent discharge by comparing them with those elicited by sinusoidal stimuli. We tested whether low frequency discharge in FAII afferents evoked a clear identifiable percept of frequency and whether it was analogous to that evoked by sinusoidal stimuli within flutter range that primarily activates FAI afferents (the non-Pacinian or RA channel). Finally we evaluated frequency discrimination capacity mediated by FAII afferents (the Pacinian channel).

## Results

### Detection thresholds mediated by FAII afferents at low frequencies

Detection thresholds for pulsatile stimuli (Fig. 1b) evoking low frequency discharge exclusively in FAII afferents (the Pacinian channel) were measured at two frequencies within the flutter range (6 and 24 Hz) and for comparison at two frequencies in the vibratory hum range (100 and 200 Hz). For pulsatile stimuli, the detection thresholds on the finger were low at all frequencies: 1.3 (±0.6 mean±SD; n=6) μm at the lowest (6 Hz) frequency and 0.7 (±0.2; n=6) μm at the highest (200 Hz) (Fig. 1a). Regardless of the frequency, the perceptual thresholds for pulsatile stimuli were well below response threshold for FAI afferents (Johansson, Landstrom, & Lundstrom, 1982; H. P. Saal, Delhaye, Rayhaun, & Bensmaia, 2017; Talbot et al., 1968), and thus could only have been mediated by the FAII afferents through the Pacinian channel. The detection thresholds for sinusoidal stimuli (Fig. 1b) were considerably higher within flutter range frequencies and, as expected, steeply decreased with increasing frequency from 28 (±6; n=6) μm at 6 Hz to 0.7 (±0.2; n=6) μm at 200 Hz (Fig. 1a). This reflects a shift from activation of FAI afferents, which have thresholds around 10-15 μm even at their characteristic frequencies, to activation of the much more sensitive FAII afferents which have thresholds for sinusoidal stimulation below 1μm at their characteristic frequencies (Johansson et al., 1982; H. P. Saal et al., 2017; Talbot et al., 1968).

**Figure 1.**
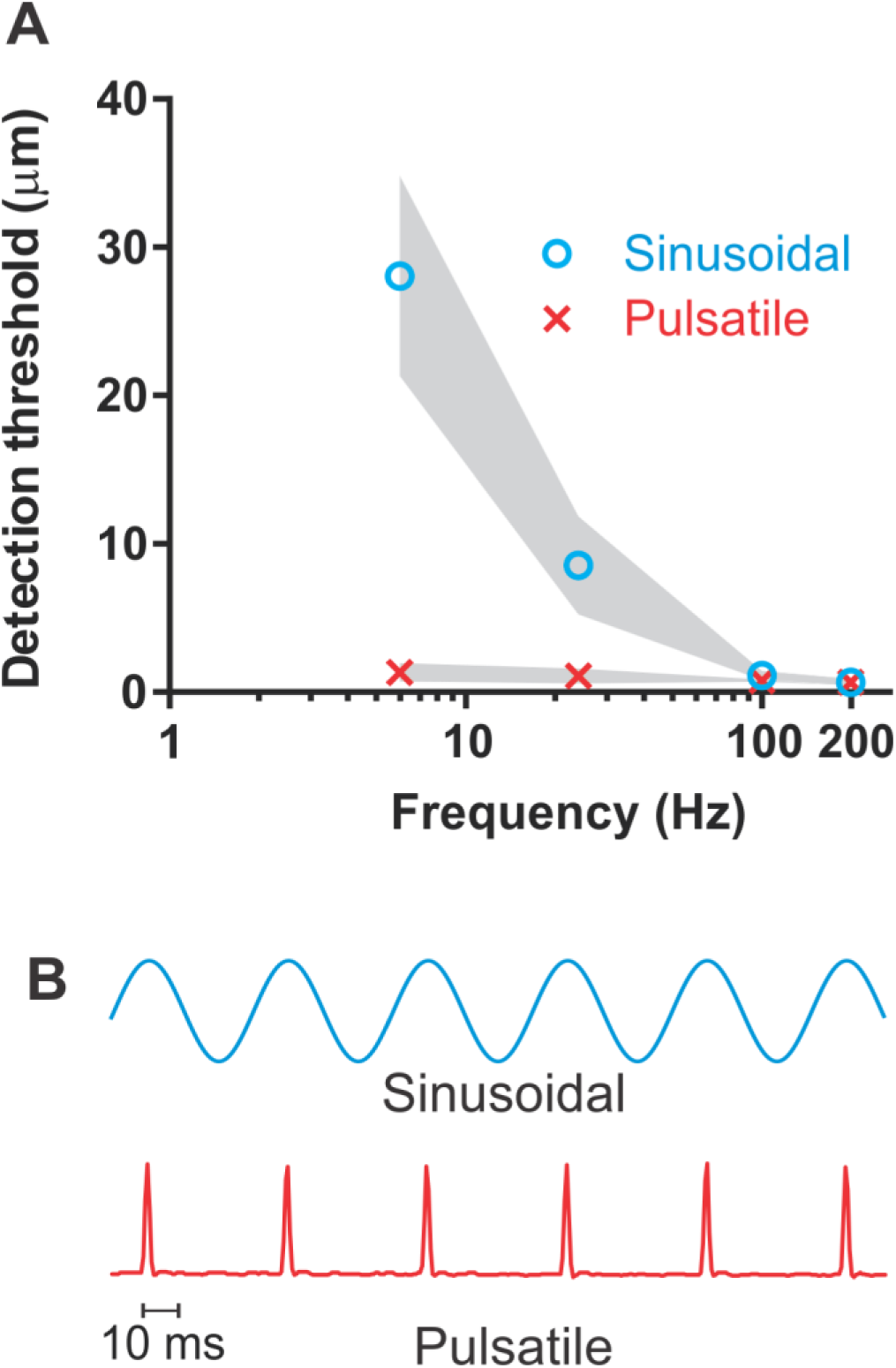
Detection thresholds. **(A)** Vibrotactile detection thresholds on the finger across frequency ranges for sinusoidal and pulsatile stimuli (n=12). Shaded area represent +/-95% confidence intervals. **(B)** An example of the sinusoidal and pulsatile waveforms. Figure 1A–Source Data 1.xlsx

### Perceptual properties of low frequency discharge rate in Pacinian channel

We next examined whether the Pacinian channel is capable of conveying a sense of vibration frequency within the flutter range. To do this, the amplitude of the pulsatile vibrotactile stimuli was kept at the level of 3 µm regardless of repetition rate (frequency), which is well below the activation thresholds of FAI afferents. The apparent frequency of this FAII-driven stimulus was obtained from participants’ comparisons of the pulsatile (P) and sinusoidal (S) stimuli in the following combinations PP, SP, SS (Fig. 2). From these comparisons, we calculated the point of subjective equality (PSE) of frequency. The physical frequency defined as repetition rate for pulsatile or frequency for sinusoidal stimuli used as test stimulus was either 20 or 40 Hz.

**Figure 2.**
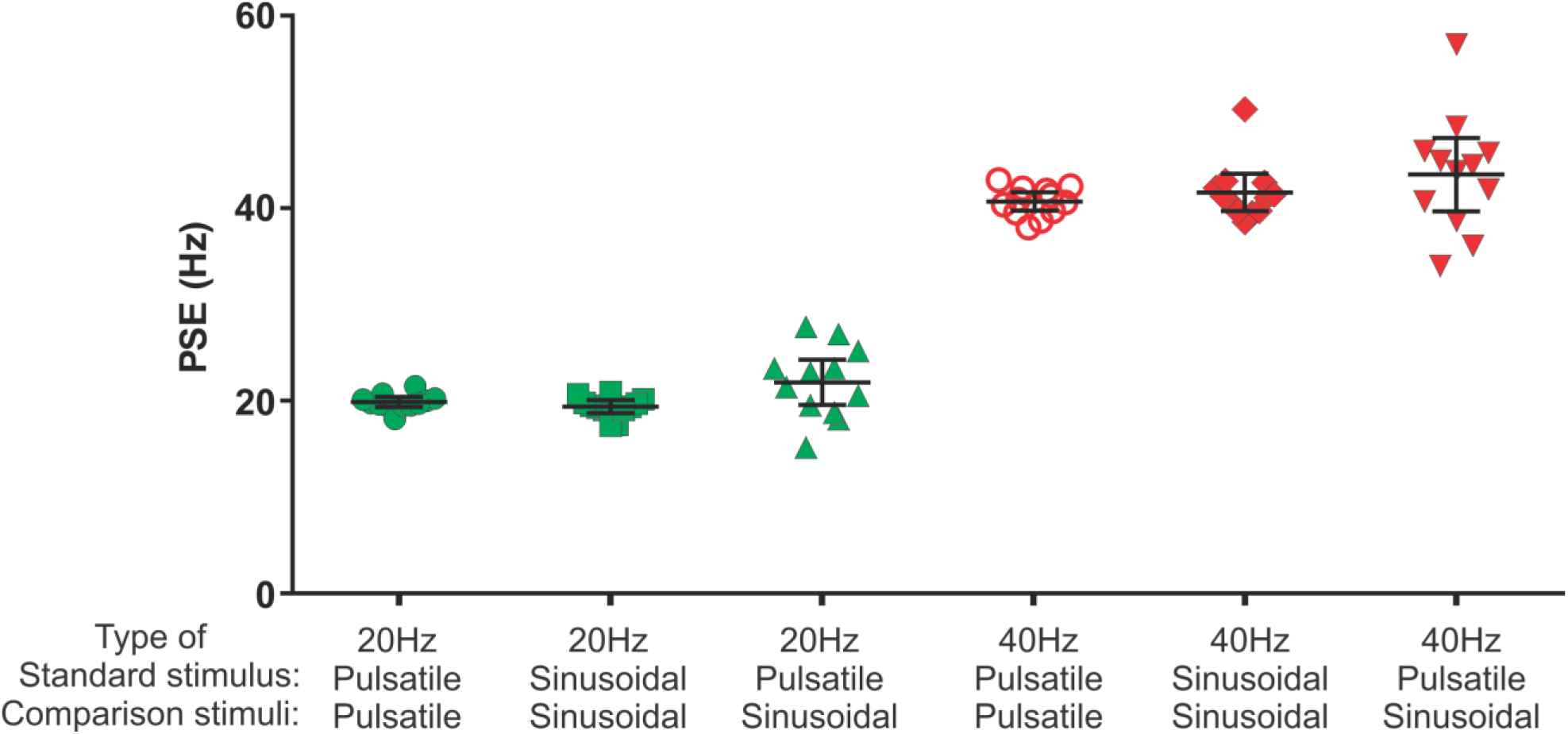
Point of subjective equality (PSE) obtained using two interval forced choice paradigm. The test stimulus was either sinusoidal or pulsatile presented at 20 Hz and 40 Hz. The test stimulus was compared with a range of comparison frequencies: 10, 14, 18, 22, 26, 30 Hz with 20 Hz test stimulus; and 25, 31, 37, 43, 49, 55 Hz with 40 Hz test stimulus. Black horizontal lines represent mean +/-95% confidence intervals (n=12). Figure 2–Source Data 1.xlsx

#### FAII afferent activation in flutter range creates frequency percept

In the PP condition, participants compared the frequency of a pulsatile test stimulus with that of six pulsatile comparison stimuli. The PSEs obtained from the psychometric curves were very close to the physical frequencies of the presented test stimuli: 20.0 (19.5 – 20.5; 95% confidence interval, CI) Hz for the 20 Hz test, and 40.8 (39.8 – 41.7) Hz for the 40 Hz test stimulus (Fig. 2; n=12). The narrow CI values indicate that pulsatile stimuli evoked perceptions with a well-defined frequency. The PSE values obtained with sinusoidal stimuli (SS condition) using six sinusoidal comparison frequencies were 19.5 (18.8 – 20.2) Hz for the 20 Hz test and 41.7 (39.8 – 43.7) Hz for the 40 Hz test stimulus (Fig. 2; n=12). Repeated measures two-way ANOVA indicated no difference between types of stimuli (pulsatile or sinusoidal) used (*F(1, 11)* = 0.131, *p* = 0.72).

#### Frequency percept mediated by low frequency discharge in FAII afferents is analogous to that evoked by sinusoidal stimuli

The PSE for the 20 Hz pulsatile test stimulus was 22.0 (19.7 – 24.4; 95% CI) Hz when determined in comparison to six sinusoidal frequencies (PS condition), which was no different from the 20 Hz stimulus (p=0.09, n=12; one sample two-tailed t-test). This indicates that both pulsatile and sinusoidal low-frequency stimuli generate a percept of identical frequency within the flutter frequency range. For 40 Hz pulsatile test stimulus, the PSE assessed in comparison with sinusoidal stimuli was 43.6 (39.8 – 47.4; 95% CI) Hz; again, this was not different from 40 Hz (p=0.06, n=12; one sample two-tailed t-test).

#### Frequency discrimination capacity mediated by FAII afferents within the flutter range

Weber fractions that were mediated exclusively by FAII afferents within the flutter frequency range were just as low as the Weber fractions determined with sinusoidal stimuli mediated predominantly by FAI afferents (Fig.3). Two-way repeated measures ANOVA indicated that the FAII afferents provided frequency discrimination in the flutter range that was no different from sinusoidal stimuli predominantly mediated by FAI afferents (*F(1, 11)* = 0.004, *p* = 0.949). However there was an effect of frequency (*F(1, 11)* = 29.00, *p* = 0.0002) indicating that the size of the Weber fraction is determined by frequency and not afferent type providing this input. Weber fractions were lower at the higher frequency (40 Hz) than they were at 20 Hz for both pulsatile and sinusoidal stimuli (0.21 vs 0.14 and 0.19 vs 0.15, respectively; n=12).

**Figure 3.**
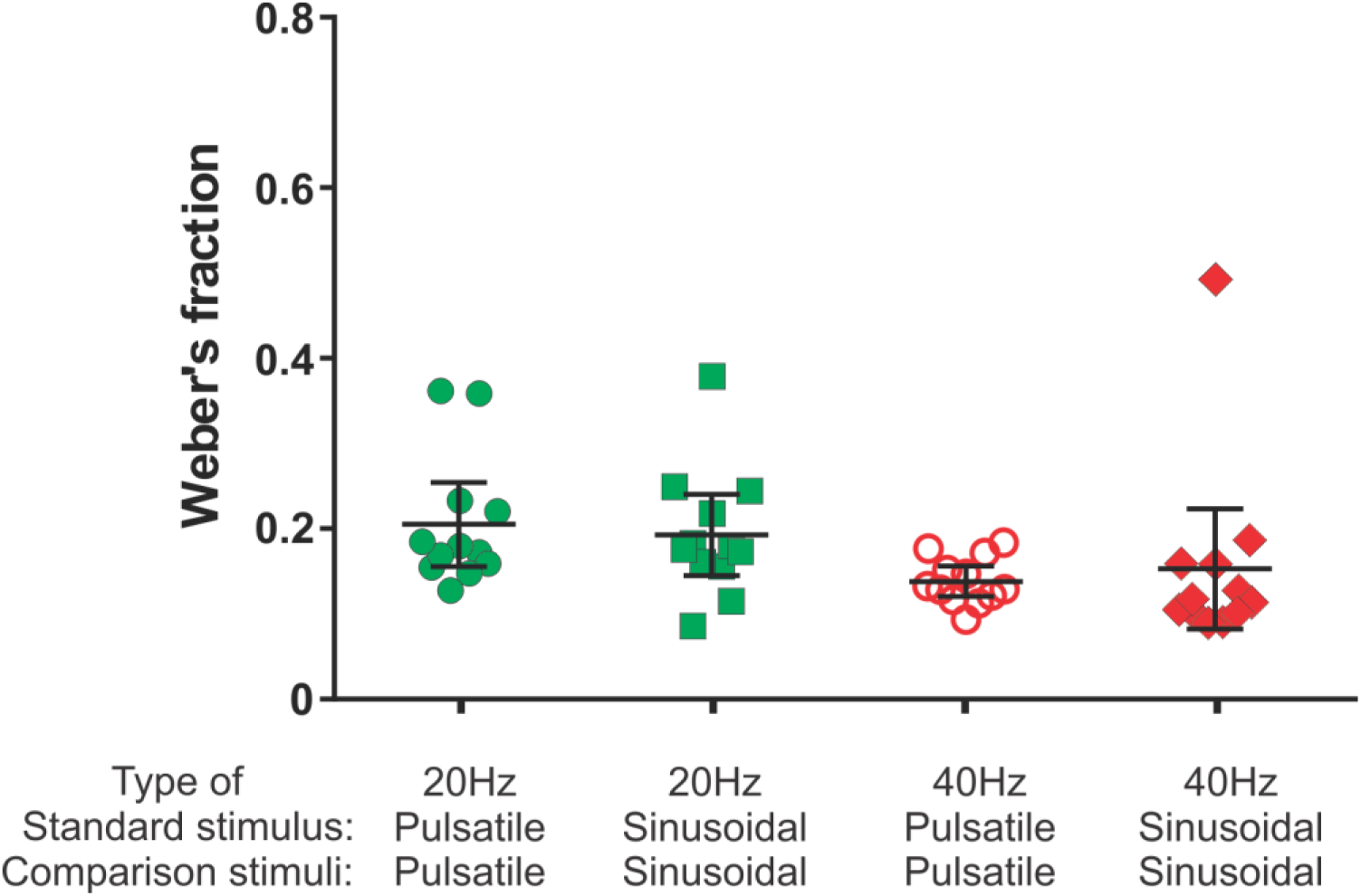
The Weber’s fraction of just noticeable difference in frequency. For details refer to legend of Figure. 2. Figure 3–Source Data 1.xlsx

## Discussion

Our study provides strong evidence that low frequency discharge of FAII afferents providing input to the Pacinian channel can mediate a clear perception of vibration with easily identifiable and distinguishable frequency characteristics within the flutter range. We also established that frequency perception signalled exclusively by the activity of FAII afferents is directly comparable with the perceptual features of flutter frequency sensation evoked by corresponding sinusoidal stimuli that naturally activate predominantly FAI afferents (non-Pacinian channel). To test whether there are inherent differences in neural mechanisms involved in frequency analysis via the Pacinian and non-Pacinian channels frequency discrimination ability was tested. The Weber fraction is generally used to characterise the smallest frequency differences reliably detected by subjects as a fraction of the comparison frequency. The Weber fraction is around 0.2 in the flutter range and at high frequencies has been reported to have a slightly higher value of around 0.3; however there is significant variation depending on methodology used (Bensmaia, Hollins, & Yau, 2005; Goble & Hollins, 1994). The discriminative ability of the Pacinian channel was not previously reported at frequencies outside the high frequency range with which it is normally associated in vibrotactile perception. The Weber fractions determined for the perceived frequency of vibration driven via the FAII and the FAI channels were a close match at both frequencies tested (20 and 40 Hz), and show a similar decrease from 20 to 40 Hz. This suggests that common neural mechanisms of frequency discrimination might be exploited by both channels, as the existence of independent mechanisms having such close agreement at frequencies outside the usual operating range of one of the channels seems less likely.

Recent psychophysical evidence demonstrated that there is interaction between the FAI and FAII inputs, by showing the assimilation effect, where a frequency in the range of one channel can influence perceived frequency on the other channel (Kuroki, Watanabe, & Nishida, 2017). We suggest that our data extends this, and represents evidence of the functional consequence of the recently discovered extensive convergence of FAI- and FAII-derived inputs onto S1 cortical neurons (Carter, Chen, Lovell, Vickery, & Morley, 2014; Pei, Denchev, Hsiao, Craig, & Bensmaia, 2009; Hannes P. Saal, Harvey, & Bensmaia, 2015). It is known that about 70-80% of rapidly-adapting neurons in somatosensory cortex S1 show convergent inputs deriving from both afferent classes, and it may be that these neurons are responsible for this generalised frequency processing. Saal et al. (Hannes P. Saal et al., 2015) made an interesting observation that input from FAI afferents determines the cortical neuron response rate due to its net excitatory drive, while the more temporally-precise PC-channel has a balanced excitatory-inhibitory drive that can control the precise spike timing, which is useful in various encoding schemes (Birznieks & Vickery, 2017; Hires, Gutnisky, Yu, O’Connor, & Svoboda, 2015; Johansson & Birznieks, 2004; Prsa & Huber, 2018; H. P. Saal, Wang, & Bensmaia, 2016). This difference did not appear to affect frequency perception in the current study as Weber fractions were found to be similar regardless of whether we used pulsatile stimuli exclusively activating FAII afferents in a time-controlled manner or used sinusoidal stimuli predominantly activating FAI afferents, presumably with less temporal precision due to the slow rising phase of the sinusoid.

A consequence of generalised neural processing for frequency, regardless of the source of afferent input, is that it would support constancy of vibrotactile frequency perception across different skin regions innervated by different afferent types. For example, the frequency perception on the hairy skin of the arm is not noticeably different from that of the glabrous skin (Mahns, Perkins, Sahai, Robinson, & Rowe, 2006; McIntyre et al., 2016), despite it having neither Meissner (FAI) nor Pacinian (FAII) receptors; instead vibrotactile stimuli are signalled by field units and hair follicle units (Vallbo, Olausson, Wessberg, & Kakuda, 1995). This also accords with natural stimulation, which is often of a sufficiently high amplitude to activate multiple types of tactile afferents (Johansson et al., 1982), and activate receptors across different skin types.

Functionally it means that FAII afferents and the Pacinian channel are well suited for detecting fast discrete mechanical transients with low repetition rate as might arise during object manipulation or exploration of surfaces with sparsely distributed sharp asperities or ridges. The evidence that low-frequency signals arising from FAII afferents are consciously perceived and easily discriminated strongly suggests that they are biologically important and are likely to be utilized by neural circuits dedicated to motor control of the hand. In regard to new technology development, the exquisite sensitivity of FAII afferents combined with their role in tactile perception and motor control makes them a useful target when designing haptic and teleoperated devices.

### Conclusions

In this study we obtained evidence that low-frequency spike trains in FAII afferents (Pacinian channel) can readily induce a vibratory percept with the same low frequency attributes as signalled by Meissner’s afferents (non-Pacinian channel). It has become evident that perception of vibrotactile frequency depends on the discharge *pattern* of the active afferents, rather than the afferent *type* that is active. Low-frequency spike trains in FAII afferents can induce a vibratory percept which has the same frequency attributes as that induced by sinusoidal stimuli. These new findings raise questions about whether much of the observed functional dichotomy between Pacinian and non-Pacinian channels relates to behavioural interpretations of the stimulus rather than to the type of receptor that the signal originates from. In addition, our proposed universal frequency decoding system would help explain the perceptual constancy of vibrotactile frequency perception which is a prominent problem in tactile system where distinct human skin regions and types (e.g., glabrous and hairy) functionally encode the same physical features of stimuli using remarkably different receptor types tuned for different stimulus features.

These findings are consistent with the growing evidence of extensive convergence of inputs from different afferent types onto neurons in the primary somatosensory cortex. Finally, these findings indicate the need to review the functional and neurophysiological basis on which processing of vibrotactile stimuli is attributed to Pacinian and non-Pacinian channels.

## Materials and Methods

### Subjects

Research participants were healthy volunteers aged 20 to 26 years without any known history of neurological disorders which would affect the somatosensory system. Ethics approval was obtained from the UNSW Human Research Ethics Committee, and all participants signed a consent form. The participants were reimbursed for their time. There were 6 participants (4 female) in the detection threshold experiments and 12 (6 female) in the frequency perception experiments. Five participants were in both experiments. The sample size was determined by pilot studies to determine effect size, and according to accepted practice in psychophysical experiments. No individual subjects or data outliers were excluded from the data analyses.

### Apparatus

The mechanical stimulation probe was a metal ball 5 mm in diameter at the end of a metal rod driven by a V4 shaker (Data Physics, San Jose, USA). To drive the shaker, analog output signals were amplified by a Signalforce™ 30W Power Amplifier (Data physics, San Jose, USA). The displacement of the stimulation probe was monitored using an OptocoNCDT 2200-10 laser displacement sensor (Micro-Epsilon, Ortenburg, Germany) with a resolution of 0.15 μm at 10 kHz.

Stimulus delivery was controlled by a CED data acquisition system (CED, Cambridge, UK) consisting of hardware (CED Power 1401 MkII) and software (Spike2 7.07). Custom made Spike2 and MATLAB (MathWorks Inc, Natick, MA, USA) scripts were used to control the delivery of pulsatile and sinusoidal stimuli, and to record stimulus measurements and button presses made by the participant.

The stimuli were delivered to the finger pad of the right index finger. The arm, hand and stimulated finger were positioned and held in place with the aid of a vacuum pillow (GermaProtec, Kristianstad, Sweden). The pillow, filled with small foam balls, was moulded around the participant’s arm, and the air was then pumped out to hold its shape. The probe was positioned on the finger with a force of 50g; the probe protracted from this rest position. White noise was delivered through headphones to eliminate auditory cues associated with the mechanical stimulator. Participants made responses by pressing buttons with the unstimulated hand.

### Vibration stimulus

A stereotyped brief pulsatile mechanical stimulus with a protraction time of only 2 ms was used to control the spiking pattern in recruited afferents (Fig. 1a). As the duration of the mechanical stimulus was comparable to the refractory period of the action potential, each mechanical stimulation event generated only a single time-controlled spike in responding afferents (Birznieks & Vickery, 2017). Each mechanical pulse is a reproducible and uniform event which ensures that the same population of afferents will be excited regardless of the rate at which these pulses are repeated.

### Detection thresholds

Detection thresholds were measured for pulsatile and sinusoidal stimuli at four frequencies: 6, 24, 100 and 200 Hz. Thresholds were determined on the fingertips. All together thresholds were tested in 16 conditions (2 waveforms x 4 frequencies x 2 locations). The thresholds for two types of stimuli were measured together in a single session with their trials pseudo-randomly interleaved. Each testing session lasted about 10 minutes, with a total of 8 sessions for each participant.

To measure detection thresholds, we used a two-interval forced-choice (2IFC) procedure, where in each trial participants were presented with two time intervals, indicated with audio cues (Fig. 4a). The intervals were each 1 s long, with a 0.5 s gap in between. One interval contained the vibration, and the other did not. Participants had to indicate which interval, the first or the second, contained the vibration stimulus. The interval containing the stimulus varied randomly, with each containing the stimulus equally often throughout the experiment.

**Figure 4.**
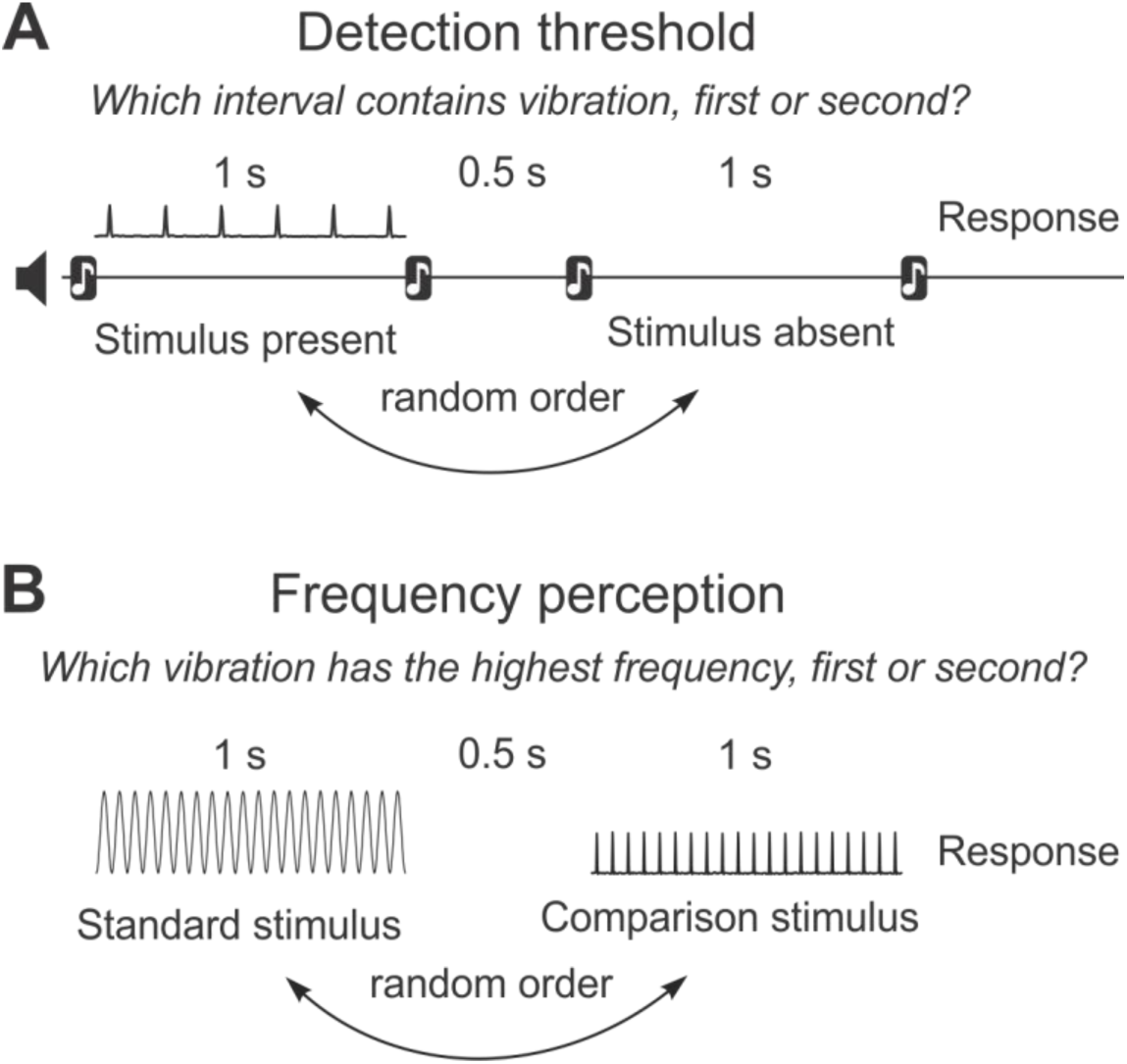
Experimental protocols. **(A)** Structure of the detection threshold task. **(B)** Structure of the frequency perception task.

To calculate detection thresholds, we used the QUEST package implemented in Psychtoolbox-3 (http://psychtoolbox.org) for MATLAB. We defined the threshold as the intensity at which the stimulus could be correctly identified for 82% of trials, and was given by the mean of the posterior distribution function. For each threshold estimate, 41 trials were conducted. To determine the amplitude of the vibration to present on each trial, we used a Bayesian adaptive QUEST protocol (Watson & Pelli, 1983), operating on the log-transformed amplitudes. The prior threshold estimate depended on the waveform and frequency (80 µm for 6 Hz sinusoidal, 5 µm for 24 Hz sinusoidal and 3 µm for 100 and 200 Hz sinusoidal, and for all pulsatile stimuli). The amplitude of the vibration on each trial was determined by the QUEST algorithm in most cases. The exceptions were the first trial, which was fixed at the prior threshold estimate for that stimulus, and every tenth trial, which was 3 times the value suggested by QUEST, to give participants a few easy trials. The actual amplitude delivered was measured and this value, along with the participant’s response, was returned to the QUEST algorithm on each trial.

### Frequency perception

To measure frequency perception, we used a similar 2IFC procedure as described above. However, in this case, both stimulus intervals presented to the participant contained vibration stimulation. The participant was required to indicate which of the two intervals contained the vibration with the highest frequency, the first or the second. One interval contained the test stimulus of fixed frequency, and the other contained the comparison stimulus, which varied in frequency from trial to trial (Fig. 4b). Six comparison frequencies were paired with the test stimulus 20 times each, in a random sequence.

Frequency perception was tested in three experimental conditions, with different combinations of stimulus waveforms: PP, in which a test stimulus with a pulsatile waveform was compared to comparison stimuli with pulsatile waveforms; SS, in which a sinusoidal test was compared to sinusoidal comparison stimuli; and SP, in which a sinusoidal test was compared to pulsatile comparison stimuli. Each condition was tested with two test stimuli of different frequencies: 20 and 40 Hz. The features of the stimuli used in the experiment are fully described in Table 1. For each condition, we randomly interleaved the trials from the 20 and 40 Hz tests. Data collecting from each participant was divided into 6 sessions lasting approximately 10 minutes each.

**Table 1.** Amplitudes and frequencies used in each experimental condition.

| Condition | Test stimulus | Comparison stimulus | Comparison frequencies |
| --- | --- | --- | --- |
| PP | 20 Hz, Pulsatile, 3 $\mu\text{m}$ | Pulsatile, 3 $\mu\text{m}$ | 10, 14, 18, 22, 26, 30 Hz |
| | 40 Hz, Pulsatile, 3 $\mu\text{m}$ | Pulsatile, 3 $\mu\text{m}$ | 25, 31, 37, 43, 49, 55 Hz |
| SS | 20 Hz, Sinusoidal, 150 $\mu\text{m}$ | Sinusoidal, 150 $\mu\text{m}$ | 10, 14, 18, 22, 26, 30 Hz |
| | 40 Hz, Sinusoidal, 40 $\mu\text{m}$ | Sinusoidal, 40 $\mu\text{m}$ | 25, 31, 37, 43, 49, 55 Hz |
| SP | 20 Hz, Sinusoidal, 150 $\mu\text{m}$ | Pulsatile, 3 $\mu\text{m}$ | 10, 14, 18, 22, 26, 30 Hz |
| | 40 Hz, Sinusoidal, 40 $\mu\text{m}$ | Pulsatile, 3 $\mu\text{m}$ | 25, 31, 37, 43, 49, 55 Hz |

The amplitude of all pulsatile stimuli was 3 µm which was approximately 3 times the sensory threshold found in our detection threshold experiment. At this amplitude, we expect only Pacinian (PC) afferents to respond, as threshold for recruitment of FAI afferents, even at their preferred frequency, is no lower than 10 µm. For the sinusoidal stimuli, we chose amplitudes such that the perceived intensity of all stimuli were approximately equal (see Table 1).

Custom MATLAB scripts were used to analyse the frequency perception data. Logistic regression was applied to the data to produce the psychometric function, relating the frequency of the comparison stimulus to the proportion of trials that the participant said the comparison was a higher frequency than the test. From the psychometric function, we calculated measures of both the perceived frequency of the test stimulus, and the frequency discrimination sensitivity. The perceived frequency of the test stimulus is given by the point of subjective equality (PSE), the comparison frequency at the 50% point on the psychometric curve (Birznieks & Vickery, 2017). The PSE is the point where the participant is equally likely to say the comparison frequency is higher or lower than the test. Discrimination sensitivity is given by the Weber fraction, the one half difference between the 75% point and the 25% on the psychometric function, divided by the frequency of the test stimulus (LaMotte & Mountcastle, 1975).

### Statistical analysis

One sample two-tailed t-test (n = 12) was used to test whether the PSE obtained in psychophysics experiments using either pulsatile or sinusoidal stimuli in twelve subjects rendered the same result as physical frequency of the periodic mechanical stimulus of the same type. In this test PSE obtained by comparing pulsatile stimulus (test stimulus) with sinusoidal stimuli (comparison stimuli) was compared to the expected PSE if test and comparison stimuli would be of the same type (sinusoidal).

Two-way repeated measures ANOVA was performed to analyse the effects on PSE by two repeated measures (within subject; n=12) factors: type of stimulus (pulsatile, sinusoidal) and frequency (20Hz, 40Hz). Two-way repeated measures ANOVA was performed to analyse the effects on Weber’s fraction (frequency discrimination capacity) by two repeated measures (within subject; n=12) factors: type of stimulus (pulsatile, sinusoidal) and frequency (20Hz, 40Hz).

For statistical analyses on the calculated thresholds, PSEs and Weber fractions, and for generating graphs, GraphPad Prism software was used (GraphPad Software Inc, La Jolla, USA).

## Acknowledgements

This work was supported by the National Health and Medical Research Council (NHMRC) project grant to IB, RMV, and VGM.

